# Potent and Selective Antitumor Activity of a T-Cell Engaging Bispecific Antibody Targeting a Membrane-Proximal Epitope of ROR1

**DOI:** 10.1101/219402

**Authors:** Junpeng Qi, Xiuling Li, Haiyong Peng, HaJeung Park, Christoph Rader

**Affiliations:** Department of Immunology and Microbiology, The Scripps Research Institute, Jupiter, Florida; X-Ray Crystallography Core, The Scripps Research Institute, Jupiter, Florida

## Abstract

T-cell engaging bispecific antibodies present a promising strategy for cancer immunotherapy and numerous bispecific formats have been developed for retargeting cytolytic T cells toward tumor cells. To explore the therapeutic utility of T-cell engaging bispecific antibodies targeting the receptor tyrosine kinase ROR1, which is expressed by tumor cells of various hematologic and solid malignancies, we used a bispecific ROR1 × CD3 scFv-Fc format based on a heterodimeric and aglycosylated Fc domain designed for extended circulatory half-life and diminished systemic T-cell activation. A diverse panel of ROR1-targeting scFv derived from immune and naïve rabbit antibody repertoires was compared in this bispecific format for target-dependent T-cell recruitment and activation. A ROR1-targeting scFv with a membrane-proximal epitope, R11, revealed potent and selective antitumor activity *in vitro* and *in vivo* and emerged as a prime candidate for further preclinical and clinical studies. To elucidate the precise location and engagement of this membrane-proximal epitope, which is conserved between human and mouse ROR1, the three-dimensional structure of scFv R11 in complex with the kringle domain of ROR1 was determined by X-ray crystallography at 1.6-Å resolution.

## Introduction

ROR1 is a receptor tyrosine kinase uniformly expressed on the cell surface of malignant B cells in chronic lymphocytic leukemia (CLL) (1-4) and mantle cell lymphoma (MCL) (5-7) but not on healthy B cells and, with some exceptions (8), other healthy cells and tissues of cancer patients. In addition to leukemia and lymphoma, ROR1 is also expressed in subsets of carcinoma, sarcoma, and melanoma, i.e., in all major cancer categories (9). Thus, ROR1 is a highly attractive candidate for targeted cancer therapy. Ongoing phase 1 clinical trials assess the safety and activity of a monoclonal antibody (mAb) (ClinicalTrials.gov identifier NCT02222688) and chimeric antigen receptor-engineered T cells (CAR-Ts) (NCT02706392) targeting ROR1 in hematologic and solid malignancies.

Using phage display, we previously generated a panel of rabbit anti-human ROR1 mAbs from immune and naïve rabbit antibody repertoires (10,11). These mAbs were shown to bind different epitopes on ROR1 with high specificity and affinity. As CAR-Ts, they mediated selective and potent cytotoxicity toward ROR1-expressing malignant cells (11,12). One of these CAR-Ts based on rabbit anti-human ROR1 mAb R12 demonstrated safety and activity in nonhuman primates (13) and was translated to the ongoing phase 1 clinical trial.

With a mechanism of action (MOA) that is conceptually related to the MOA of CAR-Ts (14,15), T-cell engaging bispecific antibodies (biAbs) are an alternative strategy for cancer immunotherapy. Like CAR-Ts, T-cell engaging biAbs utilize the power of T cells for tumor cell eradication but, unlike CAR-Ts, are manufactured and administered similarly to conventional mAbs and are cleared from the cancer patient. The concept of retargeting cytolytic T cells toward tumor cells by T-cell engaging biAbs has had substantial clinical success with a CD19 × CD3 biAb in <u>Bi</u>specific <u>T</u>-cell <u>E</u>ngager (BiTE) format, blinatumomab, receiving FDA approval for the treatment of refractory or relapsed B-cell precursor adult lymphoblastic leukemia (ALL) in 2014 (16) and with numerous other formats to various targets and indications under investigation in clinical trials (17,18). Due to their similar MOAs, both CAR-Ts and T-cell engaging biAbs can be accompanied by potentially dangerous adverse events, particularly cytokine release syndrome and central nervous system toxicity (19). Although both on-target and off-target effects contribute to adverse events, targeting cell surface antigens that are restricted to tumor cells is generally thought to afford superior safety profiles compared to targeting more widely expressed cell surface antigens.

In addition to the restricted expression of ROR1 on tumor cells, its relatively large extracellular domain (ECD) consisting of an immunoglobulin (Ig), a frizzled (Fz), and a kringle (Kr) domain, together comprising ˜375 extracellular amino acids, provides an opportunity to compare the therapeutic utility of T-cell engaging biAbs that bind to membrane-distal and membrane-proximal epitopes of ROR1. Efficient formation of cytolytic synapses (20) between T cells and tumor cells is thought to entail an optimal spacing between T-cell and tumor-cell membranes. This in turn depends on the spacing between the ROR1- and the CD3-engaging arm of the biAb as well as on the location of its epitopes on ROR1 and CD3. With a panel of rabbit anti-human ROR1 mAbs and an established humanized mouse anti-human CD3 mAb at hand, we sought to identify the best candidate for a heterodimeric and aglycosylated scFv-Fc format designed for extended circulatory half-life and diminished systemic T-cell activation. Rabbit mAb R11, which binds a conserved membrane proximal epitope on human and mouse ROR1 with similar affinity (10), emerged as the most potent ROR1-engaging arm. Its precise location and engagement was determined by co-crystallization of scFv R11 with the Kr domain of human ROR1. Collectively, our study encourages and enables the development of R11-based or R11 epitope-based T-cell engaging biAbs and other T-cell engaging cancer therapeutics.

## Results

### Design, generation, and validation of ROR1 × CD3 biAbs

Utilizing the established knobs-into-holes technology (21,22), we designed IgG1-mimicking ROR1 × CD3 biAbs in scFv-Fc format. These comprised (i) the variable light (V_L_) and heavy (V_H_) chain domains of a rabbit anti-human ROR1 mAb linked with a (Gly_4_Ser)_3_ polypeptide linker, (ii) (Gly_4_Ser)_3_-linked V_H_ and V_L_ domains of humanized mouse anti-human CD3 mAb v9 (23), which was derived from mouse anti-human CD3 mAb UCHT1 (24), and (iii) a human IgG1 Fc module with knobs-into-holes mutations for heterodimerization and with an Asn297Ala mutation to render the biAb aglycosylated (25) (**Fig. 1A**). To confirm preferential heterodimerization, we combined the scFv-Fc knobs cassette encoding vector with a vector encoding the complementary Fc holes cassette without scFv module. Analysis by SDS-PAGE revealed that >90 % of Protein A-purified antibody assembled as heterodimer (data not shown). Based on 7 rabbit anti-human ROR1 mAbs with diverse epitope specificity and affinity (10,11) (**Supplementary Table S1**), a series of ROR1 × CD3 biAbs was constructed that included R12 × v9, R11 × v9, XBR1-402 × v9, ERR1-403 × v9, ERR1-TOP43 × v9, ERR1-306 × v9, and ERR1-324 × v9. The two polypeptide chains of the heterodimeric scFv-Fc were encoded on two separate mammalian cell expression vectors based on pCEP4 and combined through transient co-transfection into Human Embryonic Kidney (HEK) 293 Phoenix cells. Non-reducing and reducing SDS-PAGE revealed the expected ˜100-kDa and ˜50-kDa bands, respectively, after purification by Protein A affinity chromatography with a yield of ˜10 mg/L (**Fig. 1B** and **Supplementary Fig. S1A**). Further analysis by size exclusion chromatography (SEC) revealed that the 7 ROR1 × CD3 biAbs eluted mainly as a single peak with <10 % aggregates (**Fig. 1C** and **Supplementary Fig. S1B**).

**Fig. 1.**
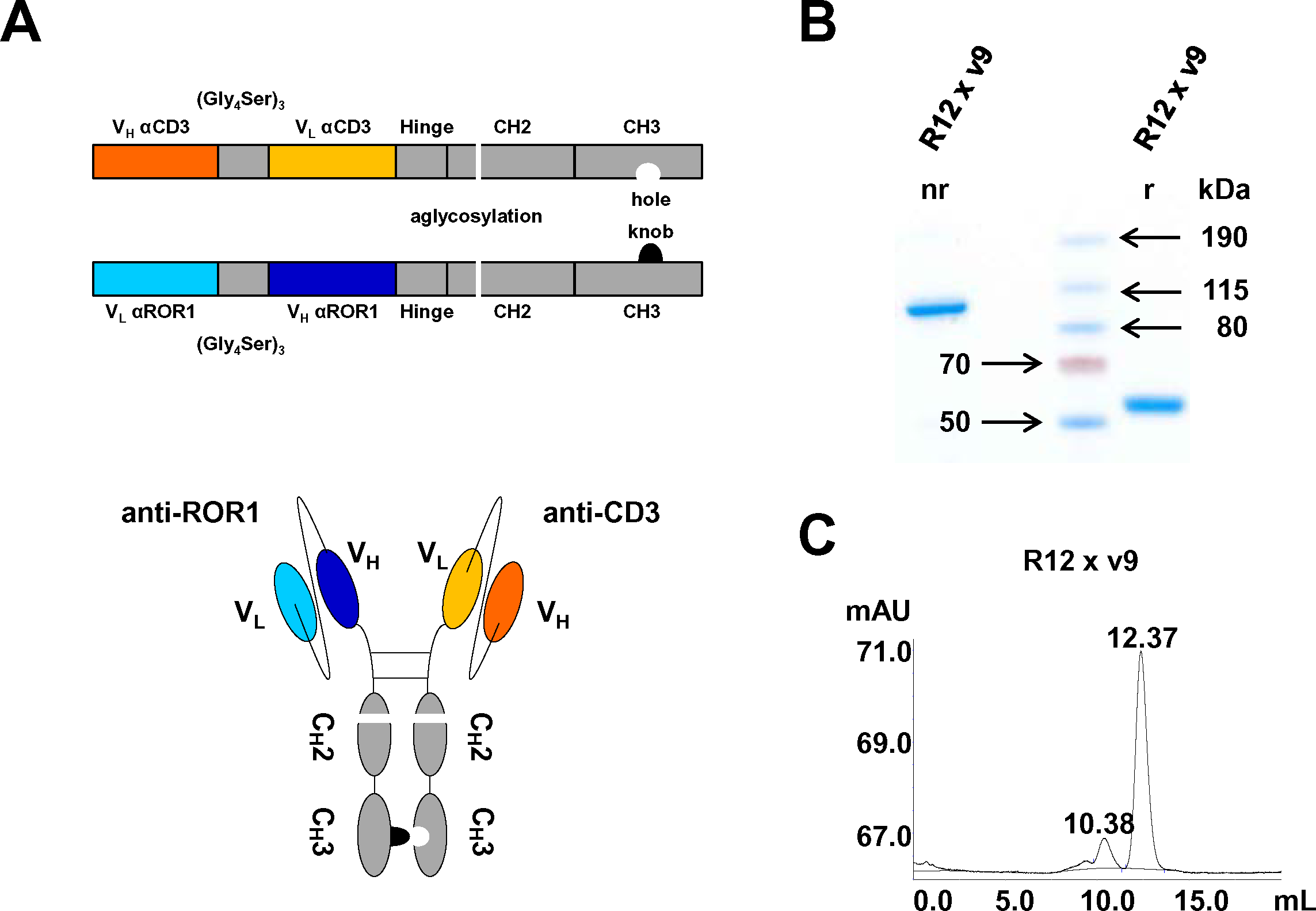
**Design and generation of bispecific ROR1 × CD3 scFv-Fc.** (**A**) Schematic diagram of ROR1 × CD3 biAb in scFv-Fc format combining a rabbit anti-human ROR1 scFv with the humanized mouse anti-human CD3 scFv v9 via a mutated Fc domain of human IgG1. The Fc domain contains 1 mutation for aglycosylation in the C_H_2 domain (N297A) and 2 or 4 mutations in the C_H_3 knob (S354C and T366W) and C_H_3 hole (Y349C, T366S, L368A, and Y407V) domains, respectively, for heterodimerization. (**B**) SDS-PAGE and Coomassie Blue staining analysis of purified representative R12 × v9 scFv-Fc showing the expected bands at ˜100 kDa under nonreducing (nr) and ˜50 kDa under reducing (r) conditions. (**C**) SEC analysis of R12 × v9 scFv-Fc eluting as major peak at 12.37 mL. The 10.38-mL minor higher molecular weight peak indicates the level of aggregation.

To study the dual binding specificity of ROR1 × CD3 biAbs, a flow cytometry assay was performed using CD3-expressing human T-cell line Jurkat, a stable K562 cell line ectopically expressing ROR1 (K562/ROR1), and MCL cell line JeKo-1 which expresses ROR1 endogenously. As shown in **Fig. 2** and consistent with the pairing of the same CD3-engaging arm with 7 different ROR1-engaging arms, all ROR1 × CD3 biAbs revealed similar binding to Jurkat cells but different binding to K562/ROR1 and JeKo-1 cells by flow cytometry. No binding to ROR1-negative parental K562 cells was detected for any of the biAbs (**Fig. 2**). To further confirm the dual binding specificity of the biAbs, we also cloned, expressed, and purified the corresponding 8 homodimeric anti-ROR1 and anti-CD3 scFv-Fc. Flow cytometry analysis of R12, R11, XBR1-402, ERR1-403, ERR1-TOP43, ERR1-306, and ERR1-324 scFv-Fc showed binding to JeKo-1 but not Jurkat cells (**Supplementary Fig. S2**). By contrast, the v9 scFv-Fc only bound to Jurkat cells (**Supplementary Fig. S2**). These data suggest that the ROR1 × CD3 biAbs retained the integrity and specificity of the parental mAbs.

**Fig. 2.**
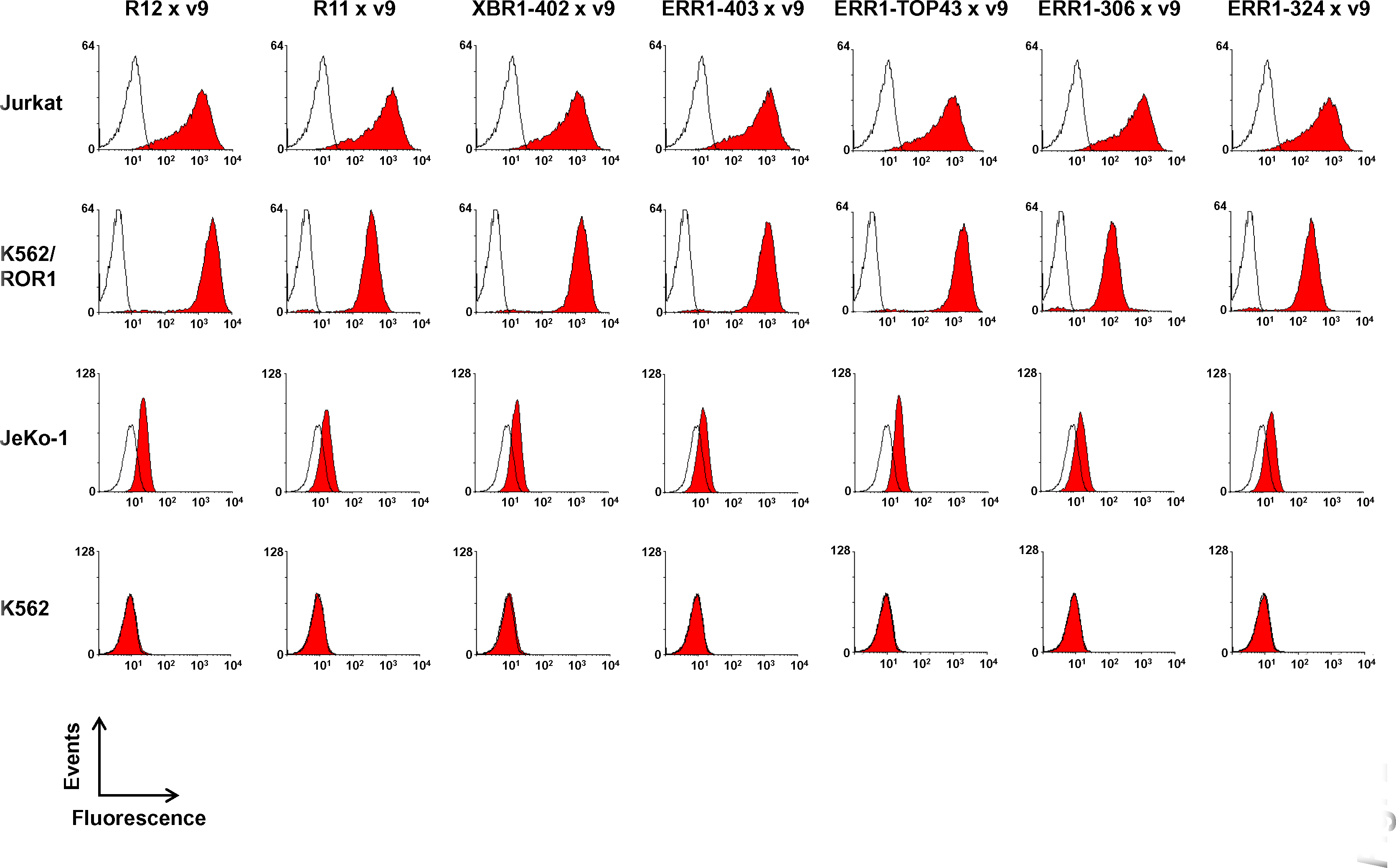
Cell surface binding of bispecific ROR1 × CD3 scFv-Fc. The indicated bispecific ROR1 × CD3 scFv were analyzed for binding to human CD3-positive T-cell line Jurkat and human ROR1-positive cell lines K562/ROR1 and JeKo-1 by flow cytometry at a concentration of 5 μg/mL followed by APC-conjugated goat anti-human IgG pAbs. Parental K562 is a ROR1-negative control cell line. Open histograms show the background binding signal of the secondary pAbs.

### *In vitro* T-cell recruitment and activation mediated by ROR1 × CD3 biAbs

We next examined the functionality of the 7 ROR1 × CD3 biAbs in terms of target-dependent T-cell recruitment and activation *in vitro*. Primary human T cells were isolated and expanded from healthy donor peripheral blood mononuclear cells (PBMC) using anti-CD3/anti-CD28 beads. The *ex vivo* expanded human T cells served as effector cells (E) and JeKo-1, K562/ROR1, and parental K562 cells as target cells (T). To study the ability of ROR1 × CD3 biAbs to recruit effector cells to target cells, we examined their crosslinking. T cells were stained red and mixed with green stained K562/ROR1 or K562 cells in the presence of 1 μg/mL ROR1 × CD3 biAbs for 1 h, gently washed, and fixed with paraformaldehyde. Double stained events detected and quantified by flow cytometry indicated crosslinked cell aggregates. As shown in **Fig. 3A** and **B**, all 7 ROR1 × CD3 biAbs demonstrated highly efficient crosslinking (˜50 %) between T cells and K562/ROR1 cells. A negative control scFv-Fc in which the ROR1-engaging arm was replaced with catalytic mAb h38C2 (26), which does not have a natural antigen, revealed only background crosslinking (<10 %). Likewise, only background crosslinking was observed when using ROR1-negative parental K562 cells (**Supplementary Fig. S3**).

**Fig. 3.**
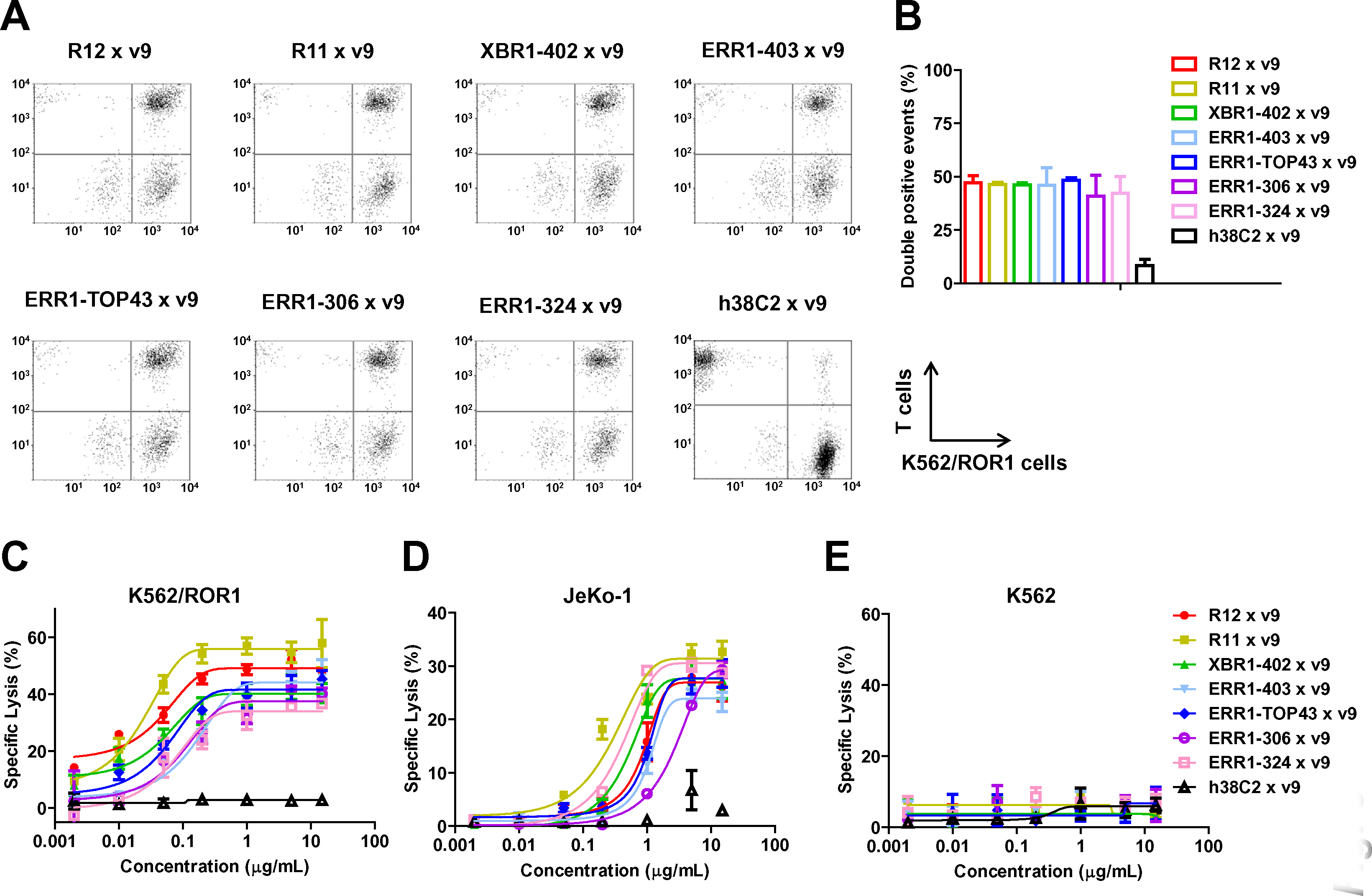
T-cell engagement mediated by bispecific ROR1 × CD3 scFv-Fc. (**A**) Crosslinking of 5 × 10^4^ primary human T cells (stained with CellTrace Far Red) and 5 × 10^4^ K562/ROR1 cells (stained with CellTrace CFSE) in the presence of 1 µg/mL ROR1 × CD3 and control scFv-Fc. Double stained events were detected by flow cytometry. (**B**) Quantification of double stained events from 3 independent triplicates (mean ± SD). (**C**) The cytotoxicity of ROR1 × CD3 scFv-Fc was tested with *ex vivo* expanded primary human T cells and K562/ROR1 (**C**), JeKo-1 (**D**), or K562 (**E**) cells at an E:T ratio of 10:1 and measured after 16 h. Shown are mean ± SD values from 3 independent triplicates.

To analyze the capability of the ROR1 × CD3 biAbs in mediating cytotoxicity *in vitro*, specific lysis of target cells after 16-h incubation with concentrations ranging from 2 pg/mL to 15 μg/mL at an E:T ratio of 10:1 was assessed. K562/ROR1 and JeKo-1 cells but not K562 cells were killed in the presence of all 7 ROR1 × CD3 biAbs, indicating that target cell lysis was strictly dependent on ROR1 (**Fig. 3C-E**). All biAbs had higher activity against K562/ROR1 cells, which express substantially higher levels of ROR1 compared to JeKo-1 cells (**Fig. 2**). Negative control biAb, h38C2 × v9, was inactive. One ROR1 × CD3 biAb, R11 × v9, was significantly and consistently more potent than the other ROR1 × CD3 biAbs and killed K562/ROR1and JeKo-1 cells with EC_50_ values of 22 ng/mL (0.2 nM) and 209 ng/mL (2 nM), respectively (**Fig. 3C** and **D**).

We next studied the ROR1 × CD3 biAbs for inducing T-cell activation. The biAbs only induced T-cell activation in the presence of K562/ROR1 cells but not K562 cells (**Fig. 4**). As analyzed by flow cytometry, the early T-cell activation marker CD69 was upregulated after 16-h incubation with 1 μg/mL ROR1 × CD3 biAbs compared to negative control biAb, h38C2 × v9 (**Fig. 4A**). As analyzed by ELISA, the release of type 1 cytokines IFN-γ, TNF-α, and IL-2 was also strictly dependent on the presence of ROR1-expressing target cells and ROR1 × CD3 biAbs (**Fig. 4B-D**). Whereas all 7 ROR1 × CD3 biAbs induced high IFN-γ secretion, R11 × v9 induced significantly higher levels of TNF-α and IL-2 secretion.

**Fig. 4.**
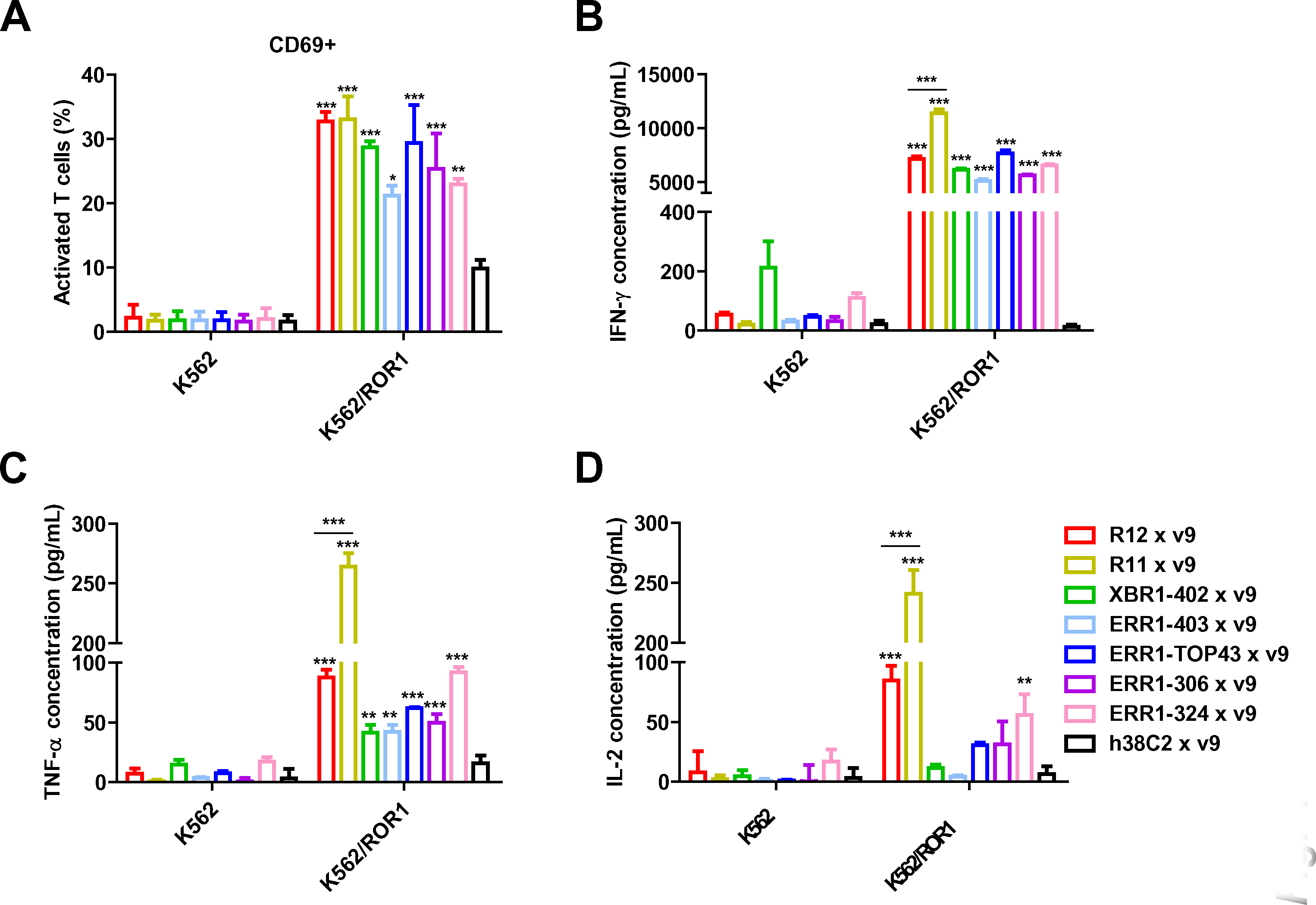
T-cell activation mediated by bispecific ROR1 × CD3 scFv-Fc. *Ex vivo* expanded primary human T cells were incubated with 1 ug/mL of the indicated biAbs in the presence of K562/ROR1 or K562 cells at an E:T ratio of 10:1 for 16 hours. (**A**) Percentage of activated T cells based on CD69 expression. Cytokine release measured by ELISA for IFN-y (**B**), TNF-a (**C**) and IL-2 (**D**). Shown are mean ± SD values for independent triplicates. One-way ANOVA was used to analyze significant differences between ROR1 × CD3 (colored) and control scFv-Fc (black) (*, p < 0.05; **, p < 0.01; ***, p < 0.001).

### *In vivo* activity of ROR1 × CD3 biAbs

To investigate whether the *in vitro* T-cell recruitment and activation mediated by ROR1 × CD3 biAbs would translate into *in vivo* activity, we used a xenograft mouse model of systemic human MCL. For this, 0.5 × 10^6^ JeKo-1 cells stably transfected with firefly luciferase (JeKo-1/ffluc) (12) were injected i.v. (tail vein) into 5 cohorts having 8 female NOD-scid-IL2Rγ^null^ (NSG) mice per cohort and, after one week when the tumor was disseminated, mice were injected i.v. (tail vein) with 5 × 10^6^ *ex vivo* expanded human T cells. One hour later, 10 μg of the ROR1 × CD3 biAbs R11 × v9 (cohort 2) and R12 × v9 (cohort 3), 10 μg of a positive control CD19 × CD3 biAb (cohort 4), and 10 μg of a negative control HER2 × v9 biAb (cohort 5) were administered i.v. (tail vein). Cohort 1 only received vehicle (PBS). All 5 cohorts were treated with a total of 3 doses of T cells (every 8 days) and 6 doses of biAbs (every 4 days). Mice were pre-conditioned with 250 μL human serum 24 h before every dose. To assess the response to the treatment, *in vivo* bioluminescence imaging was performed prior to the first dose and then every 4 days until day 39 when the signals were saturated in the control cohorts. Cohorts 1 (no biAb) and 5 (HER2 × CD3 biAb) revealed aggressive tumor growth (**Fig. 5**). In cohort 2, which received R11 × v9, we observed significant tumor growth retardation starting on day 14 and leading to nearly complete tumor eradication after 1 dose of T cells and 2 doses of biAb comparable to the CD19 × CD3 biAb in cohort 4 (**Fig. 5A** and **C**). By contrast, R12 × v9 in cohort 3 only revealed weak activity compared with the negative control cohorts. As shown in **Fig. 5B**, no weight loss was observed during the treatment in the responding cohorts, including the R11-based ROR1 × CD3 biAb which crossreacts with mouse ROR1 (**Supplementary Table S1**). Weight loss in the non-responding or weakly responding cohorts was noticeable with increasing tumor burden. The corresponding Kaplan-Meier survival curves showed that all treatment groups except for cohort 5 (HER2 × CD3 biAb) survived significantly longer than cohort 1 (no biAb). This included both ROR1 × CD3 biAb cohorts (R11 × v9, p < 0.001; R12 × v9, p < 0.05) and the CD19 × CD3 biAb cohort (p < 0.001) (**Fig. 5D**). Although all mice with this aggressive xenograft eventually relapsed, survival was longest in the CD19 × CD3 biAb cohort. As anticipated from the *in vivo* bioluminescence imaging, mice treated with the R11-based ROR1 × CD3 biAb significantly outlived mice treated with the R12-based ROR1 × CD3 biAb (p < 0.05).

**Fig. 5.**
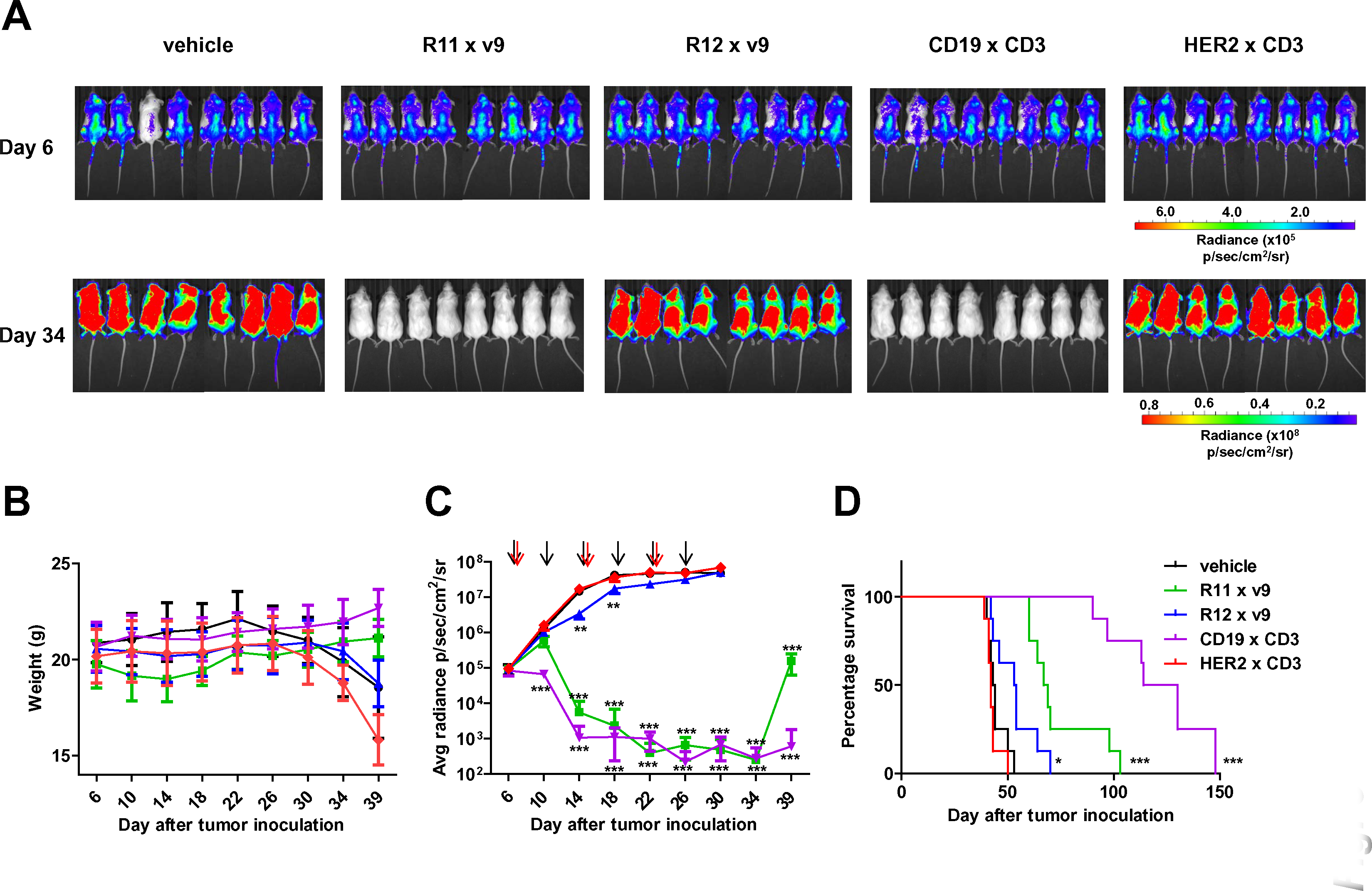
*In vivo* efficacy of bispecific ROR1 × CD3 scFv-Fc. Five cohorts of mice (n = 8) were inoculated with 0.5 × 10^6^ JeKo-1/ffluc cells via i.v. (tail vein) injection. After 7 days, 5 × 10^6^ *ex vivo* expanded primary human T cells and 10 **u**g of the indicated biAbs or vehicle alone were administered by the same route. The mice received a total of 3 doses of T cells every 8 days and a total of 6 doses of biAbs or vehicle alone every 4 days. (**A**) Bioluminescence images of all 40 mice from day 6 (before treatment) and day 34 (after treatment) are shown. (**B**) The weight of all 40 mice was recorded on the indicated days (mean ± SD). (**C**) Starting on day 6, all 40 mice were imaged every 4-5 days and their radiance was recorded (mean ± SD). Significant differences between cohorts treated with biAbs (colored graphs) and vehicle alone (black graph) were calculated using a two-tailed and heteroscedastic t-test (**, p < 0.01; ***, p < 0.001). Red arrows indicate the 3 T-cell doses and black arrows the 6 biAb or vehicle alone doses. (**D**) Corresponding Kaplan-Meier survival curves with p-values (log-rank (Mantel-Cox) test) comparing survival between cohorts treated with biAbs (colored graphs) and vehicle alone (black graph) (*, p < 0.05; ***, p < 0.001).

We next carried out a pharmacokinetic (PK) study with R11 × v9 scFv-Fc in mice to examine its circulatory half-life (t_½_). Five female CD-1 mice were injected i.v. (tail vein) with 6 mg/kg of the biAb. Blood samples were withdrawn at indicated time points (**Supplementary Fig. S4**) over a period of 2 weeks and plasma was prepared. The biAb concentration in the plasma was measured with a sandwich ELISA using immobilized ROR1 ECD for capture and peroxidase-conjugated goat anti-human IgG pAbs for detection. Analysis of the PK parameters by two-compartment modelling revealed a t_½_ of 155 ± 23 h (6.46 ± 0.96 days) (mean ± SD; **Supplementary Table S2**). For comparison, the t_½_ of both glycosylated and aglycosylated ^35^S-Met labeled chimeric mouse/human IgG1 in BALB/c mice was determined to be 6.5 ± 0.5 days (27).

### *In vitro* activity of ROR1 × CD3 biAbs toward carcinoma cell lines

To demonstrate the broader therapeutic utility of the ROR1 × CD3 biAbs beyond ROR1-expressing hematologic malignancies, we examined breast adenocarcinoma cell line MDA-MB-231 and renal cell adenocarcinoma cell line 786-O as ROR1-expressing target cells in the *in vitro* cytotoxicity assay. Prior analysis by flow cytometry confirmed that both carcinoma cell lines are recognized by all 7 ROR1 × CD3 biAbs. Among these, the R12-, XBR1-402-, and ERR1-TOP43-based ROR1 × CD3 biAbs, which are the highest affinity binders and have overlapping membrane-distal epitopes (**Supplementary Table S1**) (11), revealed the strongest binding to both carcinoma cell lines (**Fig. 6A**). Using the same cytotoxicity assay with *ex vivo* expanded human T cells as before (Fig. **3C-E**), all 7 ROR1 × CD3 biAbs revealed dose-dependent killing of MDA-MB-231 and 786-O cells (Fig. **6B** and **C**). Negative control biAb, h38C2 × v9, was again inactive. Notably, despite its weaker binding, R11 × v9 significantly outperformed all other ROR1 × CD3 biAbs for both MDA-MB-231 and 786-O cells, revealing EC_50_ values of 100 ng/mL (1 nM) and 120 ng/mL (1.2 nM), respectively (**Fig. 6B** and **C**). Collectively, these data suggest that the membrane-proximal epitope targeted by R11 is a preferred site for T-cell engaging biAbs in scFv-Fc format.

**Fig. 6.**
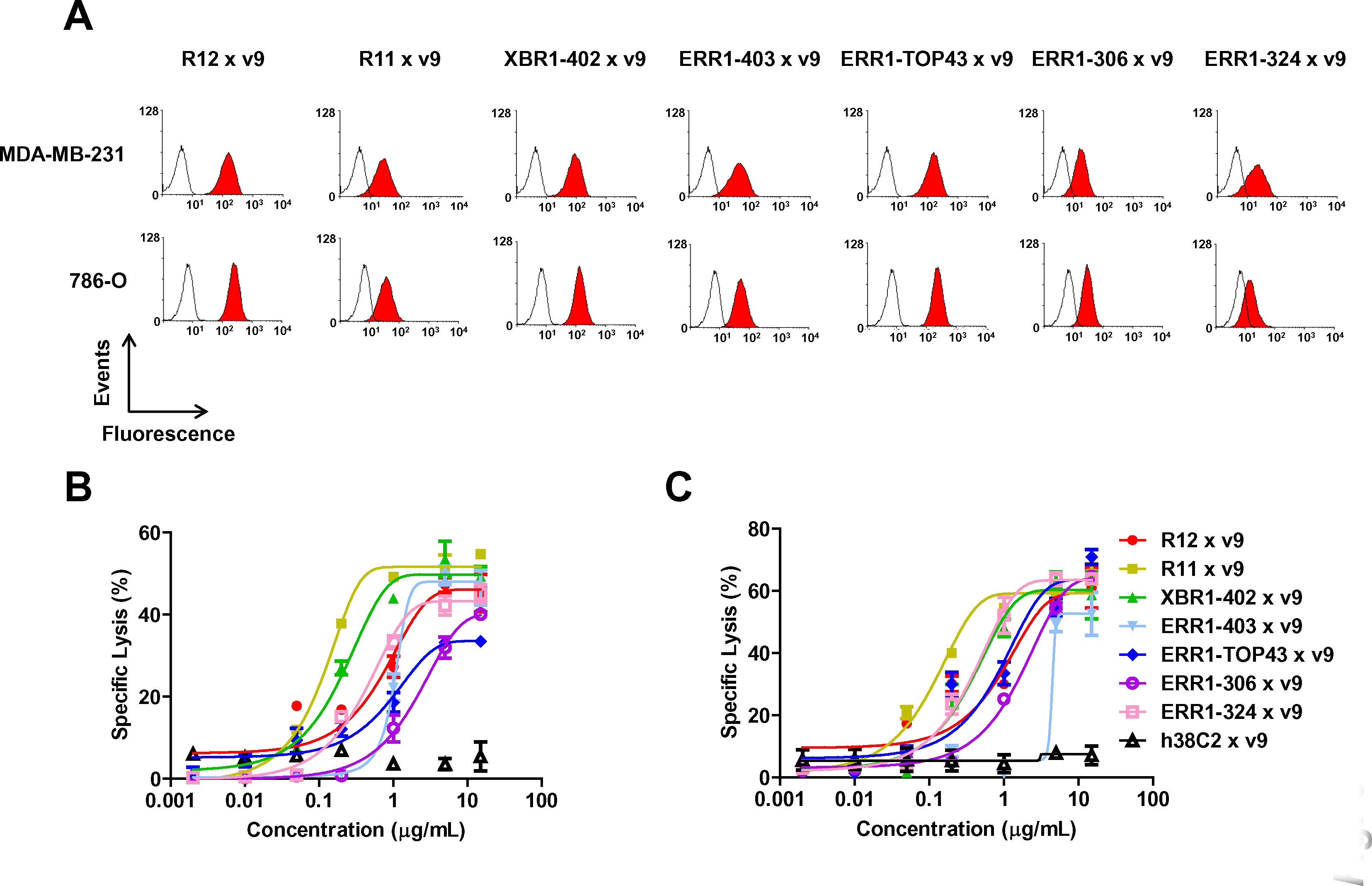
Bispecific ROR1 × CD3 scFv-Fc targeting breast cancer and renal cell cancer cell lines. (**A**) The indicated bispecific ROR1 × CD3 scFv were analyzed for binding to human ROR1-positive triple negative breast adenocarcinoma cell lines MDA-MB-231 and renal cell adenocarcinoma cell line 786-O by flow cytometry at a concentration of 5 μg/mL followed by Alexa Fluor 647-conjugated goat anti-human IgG pAbs. Open histograms show the background binding signal of the secondary pAbs. The cytotoxicity of ROR1 × CD3 scFv-Fc was tested with *ex vivo* expanded primary human T cells and MDA-MB-231 (**B**) or 786-O (**C**) cells at an E:T ratio of 10:1 and measured after 16 h. Shown are mean ± SD values from independent triplicates.

### Crystallization of R11 scFv in complex with the Kr domain of human ROR1

To elucidate the precise location of the R11 epitope, scFv R11 was crystallized in complex with the Kr domain of human ROR1 (PBD: 6BA5). The complex crystallized as P1 space group with 4 scFv:Kr complexes in the asymmetric unit (**Supplementary Table S3**). Overall the root-mean-square deviation (RMSD) of atomic positions of each complex was <0.47 Å, indicating little variation among the independent structures. The Kr domain revealed a typical kringle domain folding that appears in various extracellular proteins (**Supplementary Fig. S5**). Notably, it showed higher structural homology with the kringle domains of secreted proteins compared to the recently reported first three-dimensional structure of a kringle domain in the ECD of a transmembrane protein (28) (PDB: 5FWW). For example, the RMSD with human plasminogen kringle domain 4 (PDB: 1KRN) is only 0.66 Å. However, the shallow surface pocket known to host a free lysine or lysine analog in many kringle domains (29-31) is constricted in the ROR1 Kr domain due to an inward conformation of loop 5 and due to partial occupation by the side chain of Lys369 (**Supplementary Fig. S5**).

In the scFv:Kr complex, the *N*-terminal portion of β-strand A of the V_H_ domain undergoes domain swapping such that the first 6 amino acid integrate into the BED β-sheet of the neighboring V_H_ domain (**Fig. 7A**). The domain swapping results in crystallographic as well as non-crystallographic two-fold symmetry between two neighboring complexes and is likely a crystallization artifact as the presence of C_H_1 and C_κ_ would block this dimerization. Also, it does not influence the binding of the ROR1 Kr domain and purification by size exclusion chromatography (SEC) prior to crystallization revealed the monomeric scFv:Kr complex as major peak.

**Fig. 7.**
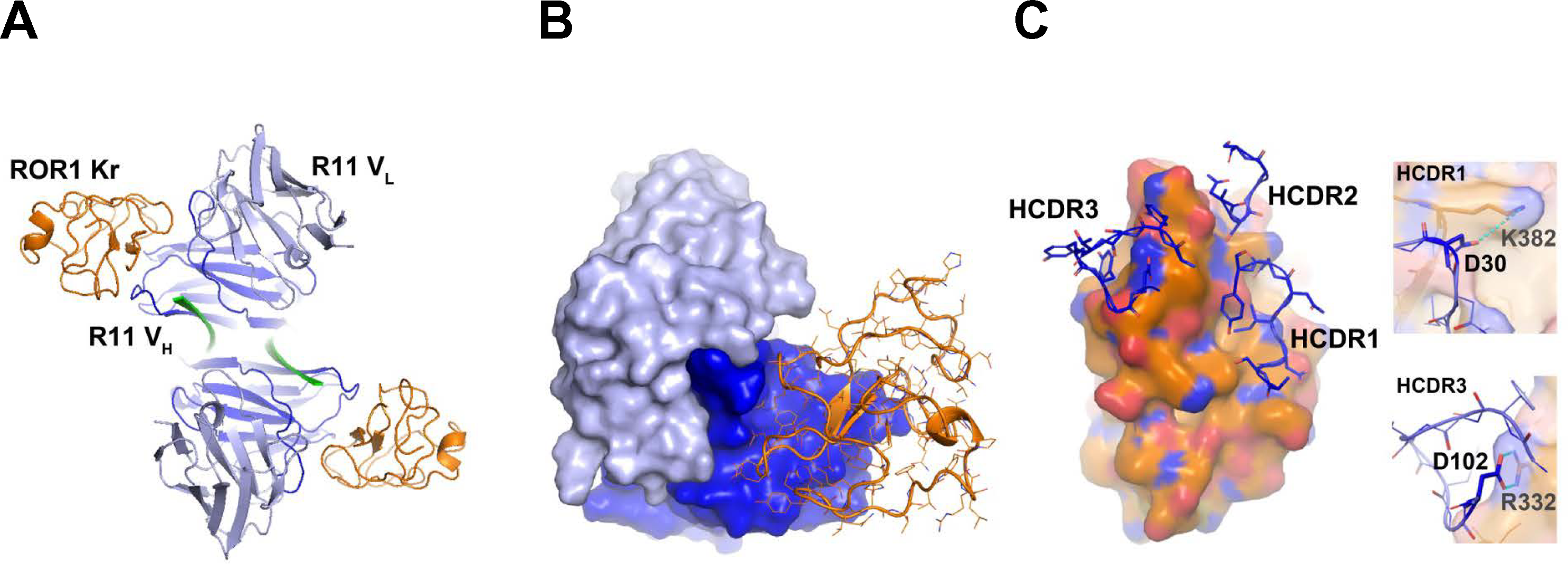
Crystal structure of scFv R11 in complex with the Kr domain of ROR1. The three-dimensional structure of scFv R11 in complex with the Kr domain of ROR1 was determined by X-ray crystallography at 1.6-Å resolution (PBD: 6BA5). (**A**) Ribbon model of the scFv:Kr complex showing two-fold non-crystallographic symmetry. The scFv is shown in dark (V_H_) and light blue (V_L_), the Kr domain in orange. The domain swapping β-strand A of the V_H_ domain is shown in green. (**B**) Surface model of the scFv showing sole interaction of the V_H_ domain (dark blue) with the Kr domain (orange tube model). (**C**) Interaction of the three CDRs of the V_H_ domain (dark blue) with the Kr domain (orange surface model). The two enlarged areas show the salt bridges formed by HCDR1 (Asp30-Lys382) and HCDR3 (Asp102-Arg332), respectively. All interacting amino acid residues are listed in Supplementary Table S4.

The total buried surface area between scFv R11 and ROR1 Kr domain is 634 Å^2^, accounting for 6 % and 13 % of the total surface areas of scFv and Kr domain, respectively. While the interaction involves multiple salt bridges, hydrogen bonds, and hydrophobic interactions, it is mediated solely by the V_H_ domain (**Fig. 7B**). No amino acid residue of the V_κ_ domain is involved in epitope recognition. Arg332 and Lys382 of the Kr domain form salt bridges with Asp102 (HCDR3) and Asp30 (HCDR1), respectively, of the V_H_ domain, enabling a strong interaction between the antibody and the antigen that is further stabilized by 9 hydrogen bonds involving all 3 CDRs of the V_H_ domain (**Supplementary Table S4**). Furthermore, multiple residues of the CDRs make van-der-Waals interactions to establish a favorable shape complementarity with the Kr domain (**Fig. 7C**). These include favorable contacts of Asp27 (HCDR1) and Tyr100 (HCDR3) with Lys369 and Arg332, respectively, of the Kr domain, which delimits the paratope, and Tyr31 and Pro32 of HCDR1 with Phe381 of the Kr domain.

The amino acid sequence identities of the Kr domains of human ROR1 vs. human ROR2 and human ROR1 vs. mouse ROR1 are 62 % and 99 %, respectively (32,33) (**Supplementary Fig. S6**). Fab R11 binds human ROR1 and mouse ROR1 with essentially the same *k*_on_, *k*_off_, and *K*_d_ (2.7 vs. 3 nM) (10), but it does not crossreact with human ROR2. **Supplementary Fig. S6** shows an amino acid sequence alignment of the 3 Kr domains with highlighted residues to indicate the epitope. These residues are highly diverse between human ROR1 and human ROR2 and completely conserved between human ROR1 and mouse ROR1, explaining the reactivity of mAb R11.

## Discussion

Harnessing and enhancing the innate and adaptive immune system to fight cancer represents one of the most promising strategies in contemporary cancer therapy. The observation that T-cell immunity plays a critical role in the immune rejection of many cancers is a key premise for cancer immunotherapy (34). A critical step of T cell-mediated eradication of tumor cells is the formation of cytolytic synapses (20) between T cells and tumor cells. This step can be mediated by biAbs combining a T-cell binding and activating arm with a tumor cell-binding arm. Numerous formats of T-cell engaging biAbs have been described and translated from basic to preclinical to clinical investigations (18). Thus far, however, only one T-cell engaging biAb, the CD19 × CD3 BiTE blinatumomab (16), has been approved by the FDA and marketed. Here we describe the generation and characterization of a panel of ROR1 × CD3 biAbs in a heterodimeric and aglycosylated scFv-Fc format that confers a circulatory half-life of nearly one week and eludes systemic T-cell activation. Built on these attributes, our panel of ROR1 × CD3 biAbs that was based on 7 different rabbit anti-human ROR1 mAbs (10,11) potently and selectively killed ROR1-expressing mantle cell lymphoma, breast cancer, and kidney cancer cells in the presence of primary T cells *in vitro*. A ROR1 × CD3 biAb that was based on a rabbit anti-human ROR1 mAb binding to a membrane-proximal epitope in the Kr domain was significantly more active *in vitro* and dramatically more active *in vivo* compared to ROR1 × CD3 biAbs with membrane-distal epitopes. Co-crystallization of the scFv in complex with the Kr domain revealed a discontinuous epitope that is conserved between human and mouse ROR1 and located in proximity to the transmembrane segment.

A recent study reported ROR1 × CD3 biAbs in BiTE format (35). The ROR1-binding arm was derived from 2 different rat anti-human ROR1 mAbs binding to an epitope in the Ig and Fz domain, respectively. The CD3-binding arm was derived from mouse anti-human CD3 mAb OKT3 (36), which shares an overlapping epitope on CD3δε with mouse anti-human CD3 mAb UCHT1 (37) and its humanized variant v9 (23,38,39) we used for our ROR1 × CD3 biAbs in scFv-Fc format. Interestingly, the BiTE targeting the Fz domain was found to be significantly more potent than the BiTE targeting the Ig domain (35). Although a three-dimensional structure of the ROR1 ECD has not been reported yet, it was assumed that the BiTE epitope in the Fz domain is in closer proximity to the cell membrane than the BiTE epitope in the Ig domain and that this closer proximity may augment the formation of cytolytic synapses between T cells and tumor cells. An increased potency of T-cell engaging biAbs that bind to membrane-proximal epitopes on tumor cell antigens has also been reported for BiTE-based MCSP × CD3 in melanoma (40) and for IgG-based FcRH5 × CD3 biAbs in multiple myeloma (41). Using confocal microscopy, the latter study provided evidence that biAbs with membrane-proximal epitopes augment the formation of cytolytic synapses by promoting target clustering and exclusion of CD45 phosphatase from the cell-cell interphase, which triggers efficient ZAP70 translocation. It was shown that this process replicates the TCR-MHC/peptide-driven formation of cytolytic synapses (41).

Epitope location on ROR1 is also critical for the activity of CAR-T cells. An R12-based CAR with membrane-distal epitope was found to be more potent with a shorter spacer between scFv and transmembrane segment whereas a longer spacer was critical for the activity of an R11-based CAR with membrane-proximal epitope (12). These findings were confirmed with an XBR1-402-based CAR, which shares an overlapping epitope with R12, and an XBR2-401-based CAR, which binds to the Kr domain of human ROR2 (11). Collectively, the screening of a panel of mAbs with diverse epitopes is critical for identifying suitable candidates for T-cell engagement by biAbs and CAR-T cells. Alternatively, if such panels are not available, customizing biAb and CAR formats may be required to incite activity. By determining the crystal structure of the R11 scFv in complex with the Kr domain of ROR1 at 1.6Å resolution, we precisely mapped the epitope to a discontinuous epitope located near the *C*-terminus of the Kr domain, 20 amino acids upstream of the predicted transmembrane segment in the primary structure of ROR1 (32). Notably, the crystal structure revealed that only V_H_ of scFv R11 makes contacts, covering ~13 % of the surface area of the Kr domain. In addition to single domain antibodies (42), our study encourages the development of R11 V_H_-mimicking non-Ig scaffolds (43), such as affibodies (44) and designed ankyrin repeat proteins (DARPins) (45), in order to explore alternative building blocks for ROR1 × CD3 engagement.

Despite being smaller, the scFv-Fc format (~100 kDa) mimics the natural IgG1 format (~150 kDa) in utilizing neonatal Fc receptor (FcRn)-mediated recycling for extended circulatory half-life. A recent study measured circulatory half-lives in mice of ~200 h for a chimeric mouse/human IgG1, ~100 h for a chimeric mouse/human scFv-Fc, and ~0.5 h for a mouse scFv, with all 3 sharing identical mouse V_L_ and V_H_ amino acid sequences (46). In approximate agreement, our PK study revealed a circulatory half-life of ~150 h for the ROR1 × CD3 scFv-Fc. This prolonged presence in the blood permitted a twice weekly (biw) dosing in our *in vivo* experiment that likely can be further reduced to once weekly (qw). By contrast, BiTEs, which are ~50-kDa (scFv)_2_ without Fc domain and a circulatory half-life of ~2 h in humans (47), require continuous administration via, e.g., a portable minipump and intravenous port catheter. While in case of blinatumomab this setting has the advantage that treatment can be interrupted to address adverse events, particularly cytokine release syndrome and neurologic events (48), an increasing number of biAb formats in preclinical and clinical studies include an Fc domain to avoid continuous intravenous infusion (18). We chose an aglycosylated Fc domain to retain FcRn-mediated recycling but elude binding to Fcγ receptors CD16, CD32, and CD64 (49). Although this design excludes antibody-dependent cellular cytotoxicity (ADCC) and antibody-dependent cellular phagocytosis (ADCP) as contributing effector functions, it ensures that T cells are not activated systemically by Fcγ receptor-bound scFv-Fc that crosslink CD3 (50). Using an aglycosylated Fc did not impact expression yields in mammalian cells, purification yields using standard Protein A affinity chromatography, monodispersity, and solubility of the scFv-Fc, affirming its suitability for manufacturing.

Despite limited insights into its role in cancer biology, the receptor tyrosine kinase ROR1 is an attractive target for antibody-mediated cancer therapy due to its highly restricted expression in a range of hematologic and solid malignancies (9). Postnatal healthy cells and tissues are largely negative for cell surface ROR1 despite some exceptions (6-8,51). Two early clinical trials currently investigate ROR1 as a target for mAb and CAR-T cell therapy, respectively. ROR1 × CD3 biAbs, which combine off-the-shelf availability of mAbs with T-cell engaging potency of CAR-T cells, could provide a best-of-both-worlds option for patients with ROR1-expressing cancers. Due to its potency *in vitro* and *in vivo*, its recent humanization (52), and the precise mapping of its paratope and epitope by X-ray crystallography, mAb R11 has emerged as a prime candidate for further preclinical and clinical studies of ROR1 × CD3 biAbs.

## Materials and Methods

### Cell lines and primary cells

The K562, JeKo-1 and Jurkat cell lines were obtained from the American Type Culture Collection (ATCC) and cultured in RPMI-1640 supplemented with L-glutamine, 100 U/mL Penicillin-Streptomycin, and 10 % (v/v) fetal calf serum (FCS; all from Thermo Fisher Scientific). The K562/ROR1 and JeKo-1/ffluc cell lines (6,12) were kindly provided by Drs. Stanley R. Riddell (The Fred Hutchinson Cancer Research Center, Seattle, WA) and Michael Hudecek (University of Würzburg, Würzburg, Germany), respectively. HEK 293 Phoenix (ATCC), MDA-MB-231 (ATCC), and 786-O cells (an NCI-60 panel cell line obtained from The Scripps Research Institute’s Cell Based High Throughput Screening Core) were grown in DMEM supplemented with L-glutamine, 100 U/mL Penicillin-Streptomycin, and 10 % (v/v) FCS. PBMC were isolated from healthy donor buffy coats using Lymphoprep (Axis-Shield) and cultured in X-VIVO 20 medium (Lonza) with 5 % (v/v) off-the-clot human serum AB (Gemini Bio-Products) and 100 U/mL IL-2 (Cell Sciences). Primary T cells were expanded from PBMC as previously described (31) using Dynabeads® ClinExVivo™ CD3/CD28 (Thermo Fisher Scientific).

### Cloning, expression, and purification of ROR1 × CD3 biAbs

All amino acid sequences have been previously published or patented. The variable domain encoding cDNA sequences of the rabbit anti-human ROR1 m Abs and the humanized mouse anti-human CD3 mAb v9 were PCR-amplified from phagemids (10,11) and previously described DART-encoding plasmids (53), respectively. A (Gly_4_Ser)_3_ linker encoding sequence was used to fuse V_L_ and V_H_ by overlap extension PCR. The Fc fragment with a N297A mutation and either knob mutations, S354C and T366W, or hole mutations, Y349C, T366S, L368A, and Y407V, were synthesized as gBlocks (Integrated DNA Technologies). Anti-ROR1-Fc knob and anti-CD3-Fc hole encoding fragments were assembled by overlap extension PCR, respectively, *Asc*I/*Xho*I-cloned into mammalian cell expression vector pCEP4, and transiently co-transfected into HEK 293 Phoenix cells (ATCC) with polyethylenimine (Sigma-Aldrich) as described (54). Supernatants were collected 3 times over a 9-day period followed by 1-mL Protein A HiTrap HP columns in conjunction with an ÄKTA FPLC instrument (both from GE Healthcare Life Sciences). Subsequent preparative and analytic (10 μg biAb) size exclusion chromatography was performed with a Superdex 200 10/300 GL column (GE Healthcare Life Sciences) in conjunction with an ÄKTA FPLC instrument using PBS at a flow rate of 0.5 mL/min. The purity of the biAbs was confirmed by SDS-PAGE followed by Coomassie Blue staining and their concentration was determined by measuring the absorbance at 280 nm.

### Flow cytometry

Flow cytometry was performed on a FACSCanto (BD Biosciences) and data were analyzed with WinMDI 2.9 software. APC or Alexa Fluor 647-conjugated goat anti-human IgG pAbs were purchased from Jackson ImmunoResearch Laboratories. Alexa Fluor 647-conjugated mouse antihuman CD69 mAb was purchased from BioLegend. For the crosslinking assay, target and effector cells were labeled with CellTrace CFSE and CellTrace Far Red (both from Thermo Fisher Scientific), respectively, according to the manufacturer’s protocol. The labeled target and effector cells at a 1:1 ratio were then incubated with 1 µg/mL biAbs in 100 µL final volume. Following incubation for 1 h at room temperature, the cells were gently washed, fixed with 1 % (w/v) paraformaldehyde, and analyzed by flow cytometry as described above.

### Cytotoxicity assay

Cytotoxicity was measured using CytoTox-Glo (Promega) following the manufacturer’s protocol with minor modifications. Primary T cells expanded from healthy donor PBMC as described above were used as effector and K562/ROR1, JeKo-1, K562, MDA-MB-231, or 786-O cells were used as target cells. The cells were incubated at an effector-to-target (E:T) ratio of 10:1 in X-VIVO 20 Medium (Lonza) with 5 % (v/v) off-the-clot human serum AB. The target cells (1 × 10^4^) were first incubated with the biAbs prior to adding the effector cells (1 × 10^5^) in a final volume of 100 µL/well in a 96-well tissue culture plate. The plates were incubated for 16 h at 37 °C with biAb concentrations ranging from 2 pg/mL to 15 µg/mL. After centrifugation, 50 µL of the supernatant was transferred into a 96-well clear bottom white walled plate (Costar 3610; Corning) containing 25 µL/well CytoTox-Glo. After 15 min at room temperature, the plate was read using a SpectraMax M5 instrument with SoftMax Pro software. The same supernatants (diluted 20-fold) used for the CytoTox-Glo assay were also used to determine IFN-γ, TNF-α, and IL-2 secretion with the Human IFN gamma, Human TNF alpha, and Human IL-2 ELISA Ready-SET-Go! reagent sets (eBioscience), respectively, following the manufacturer’s protocols.

### Mouse xenograft studies

Forty 6-weeks old NOD-scid-IL2Rγ^null^ (NSG) mice (The Jackson Laboratory) were each given 0.5 × 10^6^ JeKo-1/ffluc cells via i.v. (tail vein) injection on day 0. On day 7, each mouse was i.v. (tail vein) injected with 5 × 10^7^ primary T cells expanded from healthy donor PBMC as described above, and 1 h later with 10 µg biAbs or with PBS alone. The mice received a total of 3 doses of expanded primary T cells every 8 days and a total of 6 doses of biAbs or PBS alone every 4 days. Every 4-5 days, tumor growth was monitored by bioluminescent imaging 5 min after i.p. injections with 150 mg/kg D-luciferin (Biosynth). For this, mice were anesthetized with isoflurane and imaged using an Xenogen IVIS Imaging System (Caliper) 6, 8, 10, 12, and 14 minutes after luciferin injection in small binning mode at an acquisition time of 10 s to 1 min to obtain unsaturated images. Luciferase activity was analyzed using Living Image Software (Caliper) and the photon flux analyzed within regions of interest that encompassed the entire body of each individual mouse. The weight of the mice was measured every 4-5 days. All procedures were approved by the Institutional Animal Care and Use Committee of The Scripps Research Institute and were performed according to the NIH Guide for the Care and Use of Laboratory Animals.

### Pharmacokinetic study

Five female CD-1 mice (~25 g; Charles River Laboratories) were injected i.v. (tail vein) with R11 × v9 at 6 mg/kg. Blood was collected at 5 min and 12, 24, 48, 72, 108, 168, 240, and 336 h post-injection with heparinized capillary tubes. Plasma was obtained by centrifuging the samples at 2,000 g for 5 min in a microcentrifuge and stored at -80 °C until analysis. The concentrations of biAbs in the plasma samples were measured using ELISA. For this, each well of a 96-well Costar 3690 plate (Corning) was incubated with 200 ng recombinant human ROR1 ECD (10) in 25 μL carbonate/bicarbonate buffer (pH 9.6) at 37 °C for 1 h. After blocking with 150 μL 3 % (w/v) BSA/PBS for 1 h at 37 °C, the plasma samples were added. Peroxidase AffiniPure F(ab')_2_ Fragment Goat Anti-Human IgG pAbs (Jackson ImmunoResearch Laboratories) were used for detection. The concentration of the biAbs in the plasma samples was extrapolated from a four-variable fit of the standard curve. Pharmacokinetic parameters were analyzed using Phoenix WinNonlin PK/PD Modeling and Analysis software (Pharsight).

### Statistical analyses

Statistical analyses were performed using Prism Software (GraphPad). The *in vitro* data shown in **Fig. 4A-D** were subjected to one-way analysis of variance (ANOVA) and the *in vivo* data shown in **Fig. 5C** were analyzed with a two-tailed and heteroscedastic t-test with a confidence interval of 95 %. Statistical analysis of survival (**Fig. 5D**) was done by log-rank (Mantel-Cox) testing. Results with a p-value of p < 0.05 were considered significant.

### Crystallization and structure determination

*Cloning, expression, and purification* – Leaderless R11 scFv and human ROR1 Kr domain (amino acids 319-391) encoding cDNA sequences were PCR-amplified from previously described phagemids and plasmids (10) and cloned as polycistronic assembly that also included a leaderless *E. coli* disulfide bond isomerase (DsbC), which is a chaperone that aids proper disulfide bond formation (55), into *E. coli* expression vector pET-15b (Novagen). The resulting expression cassette included a ribosome binding site (RBS) with a start codon, an *N*-terminal (His)_6_ tag, a thrombin cleavage site, the Kr domain (flanked by *Nde*I/*Xho*I), a second RBS with a start codon followed by the scFv (flanked by *Xho*I/*Xho*I), and a third RBS with a start codon followed DsbC (flanked by *Xho*I/*Bam*HI). The prepared plasmid was transformed into *E. coli* strain Rosetta-Gami 2(DE3) (Novagen). Following induction of protein expression by 0.3 mM isopropyl-D-thiogalactoside, the bacterial cultures were grown in LB medium containing ampicillin, tetracycline, and chloramphenicol at 20 °C and 220 rpm for 18 h.

*Protein Purification* – The proteins were purified by immobilized metal ion chromatography followed by size exclusion chromatography in conjunction with an ÄKTA FPLC instrument. In brief, bacterial pellets were harvested by centrifugation and resuspended in sonication buffer (20 mM HEPES (pH 8.0), 300 mM NaCl, 15 mM imidazole, 10 % (v/v) glycerol), sonicated in an ice-water bath, and centrifuged for 25 min at 53,300 g. The supernatant was loaded on a custom-packed 10-mL HIS-Select column (Sigma-Aldrich) and washed with sonication buffer. The scFv:Kr complex was eluted with a linear gradient of 15-500 mM imidazole and treated overnight at 4 °C with thrombin (Sigma-Aldrich) to remove the *N*-terminal (His)_6_ tag on the scFv. The clipped scFv:Kr complex was then purified further by size-exclusion chromatography on a Superdex 200 26/60 column (GE Healthcare Life Sciences) equilibrated with 50 mM NaCl, 10 mM HEPES (pH 7.4). Fractions containing the complex were combined and concentrated to ~15 mg/mL with a 10-kDa MWCO Amicon ultrafiltration unit (Millipore).

*Crystallization and structure determination –* Crystals of the purified scFv:Kr complex were grown by vapor diffusion at room temperature using 1.5 µL of 15 mg/mL protein and an equal volume of precipitant containing 0.2 M ammonium fluoride in 20 % (w/v) polyethylene glycol 3,350, and were fully grown within 2 days. The crystals were flash frozen in liquid nitrogen using nylon loops after removing excess mother liquor. A diffraction data set with Bragg spacings to 1.6 Å was collected on an ADSC Quantum 315r detector at the 24-ID-E beamline at the Advanced Photon Source, Argonne National Laboratory. Data was processed with iMOSFLM software (56). The structure of the scFv:Kr complex was solved by the molecular replacement method using BALBES with 2ZNW as the search model (57). Crystallographic refinement was performed using a combination of PHENIX 1.12 (58) and BUSTER 2.9 (59). Manual rebuilding and adjustment of the structure were carried out using the graphics program Coot (60). Data processing and refinement statistics are shown in **Supplementary Table S3**. Molecular figures were created using PyMOL software (Schrödinger). Buried surface area was calculated using PISA (61) and structure validation was carried out with MolProbity (62). The crystal structure was deposited in PDB as 6BA5.

## Acknowledgments

We thank Drs. Stanley R. Riddell and Michael Hudecek for cell lines and Li Lin and Dr. Michael D. Cameron for their help with analyzing the pharmacokinetic study. This study was funded by NIH grants R01 CA181258 and UL1 TR001114, and by generous donations from the PGA National Women’s Cancer Awareness Days, the Jewish Federation of Palm Beach County, and the Anbinder Family Foundation. This is manuscript 29607 from The Scripps Research Institute.

## Supplementary Tables

**Supplementary Table S1.** Properties of rabbit anti-human ROR1 mAbs used in this study.

|  | **R12** | **R11** | **XBR1-402** | **ERR1-403** | **ERR1-TOP43** | **ERR1-306** | **ERR1-324** |
| --- | --- | --- | --- | --- | --- | --- | --- |
| **HCDR3** | DSYADDGALFNI | GYSTYYGDFNI | DHPTYGMDL | GSDYFDL | DVHSTATDL | WVYGVDDYGDGNWLDL | GTVSSDI |
| **LCDR3** | GADYIGGYV | LGGVGNVSYRTS | QVWDSSAYV | QLWDSSAGAYV | QLWDSSAGAYV | GTDYSGGYV | LGGYVSQSYRAA |
| **Isotype** | $\lambda$ | $\kappa$ | $\lambda$ | $\lambda$ | $\lambda$ | $\lambda$ | $\kappa$ |
| **$K_d$ (nM)** | 0.56 | 2.7 | 5.81 | 41.6 | 1.11 | 9.66 | 108 |
| **Epitope** | Ig+Fz | Kr | Ig+Fz | Ig+Fz | Ig+Fz | Ig+Fz | Ig+Fz<sup>1</sup> |
| **hROR1<sup>2</sup>** | + | + | + | + | + | + | + |
| **cROR1<sup>3</sup>** | + | + | + | + | + | + | + |
| **mROR1<sup>4</sup>** | – | + | – | – | – | + | – |

<sup>1</sup>Non-overlapping epitope with R12, XBR1-402, ERR1-403, ERR1-TOP43, and ERR1-306
<sup>2</sup>hROR1, human ROR1; <sup>3</sup>cROR1, cynomolgus ROR1; <sup>4</sup>mROR1, mouse ROR1 reactivity.

**Supplementary Table S2.** Pharmacokinetic parameters of R11 × v9 scFv-Fc.^1^

| **t<sub>1/2</sub> (h)** | **AUC (mg/mL × h)** | **CL (mL/h/kg)** | **V<sub>ss</sub> (mL/kg)** |
| --- | --- | --- | --- |
| 155 ± 23 | 13 ± 0.8 | 0.1 ± 0.01 | 81 ± 6 |

<sup>1</sup>Mean ± SD values from 5 mice.

**Supplementary Table S3.** Data collection and refinement statistics for the crystallization of scFv R11 in complex with the Kr domain of ROR1.

|  |  |
| --- | --- |
| **Data** |  |
| Space group | P1 |
| Cell dimensions a, b, c (Å) | 58.02, 73.90, 84.12 |
| Cell dimension $\alpha, \beta, \gamma$ (°) | 89.86, 71.30, 84.49 |
| Asymmetric unit | 4 complexes |
| Resolution (Å) | 54.85 – 1.62 (1.65 – 1.62) |
| Unique reflections | 160894 (7849) |
| Mean $I/\sigma I$<sup>1</sup> | 8.9 (2.5) |
| Completeness (%)<sup>1</sup> | 96.1 (94.6) |
| Wilson B-factor | 10.18 |
| $R_{merge}$ | 0.087 (0.49) |
| $R_{meas}$ | 0.103 (0.59) |
| $R_{pim}$ | 0.055 (0.32) |
| **Refinement** |  |
| Resolution (Å) | 39.53 – 1.62 (1.66 – 1.62) |
| Reflections in refinement | 160881 (11498) |
| Reflections in $R_{free}$ | 3234 (240) |
| $R_{free}$<sup>1,2</sup> | 0.194 (0.214) |
| $R_{cryst}$<sup>1</sup> | 0.194 (0.205) |
| R.m.s. bond length (Å) | 0.01 |
| R.m.s. bond angle (°) | 1.0 |
| B-factor, average (Å<sup>2</sup>) | 22.08 |
| **Number of atoms** |  |
| Protein | 9266 |
| Water | 1364 |
| **Model Quality** |  |
| Ramachandran favored (%) | 96.74 |
| Ramachandran allowed (%) | 3.26 |
| Ramachandran outliers (%) | 0 |
| Rotamer outliers (%) | 0 |
| Clashscore | 2.15 |

<sup>1</sup>Values in parentheses refer to statistics for the highest resolution shell.
<sup>2</sup> $R_{free}$ is calculated with removal of 2 % of the data as the test set before the refinement.

**Supplementary Table S4.** List of interacting amino acid residues at the interface of scFv R11 and the kringle domain of ROR1.

|  | scFv | Type of interaction | Kr domain |
| --- | --- | --- | --- |
| **HCDR1** | Asp27 (O) | hydrogen bond | Lys369 (NZ) |
|  | Asp30 (OD1) | salt bridge | Lys382 (NZ) |
|  | Asp30 (OD1) | hydrogen bond | Ser383 (OG) |
|  | Tyr31 (OH) | hydrogen bond | Asp384 (OD1) |
| **HCDR2** | Asn51 (ND2) | hydrogen bond | Asn380 (OD1) |
|  | Ser52 (N) | hydrogen bond | Asn380 (O) |
|  | Ser52 (OG) | hydrogen bond | Lys382 (N) |
| **HCDR3** | Tyr96 (OH) | hydrogen bond | Asp378 (OD2) |
|  | Gly101 (N) | hydrogen bond | Ser330 (O) |
|  | Asp102 (OD1) | hydrogen bond | Arg332 (NH2) |
|  | Asp102 (OD2) | salt bridge | Arg332 (NH2) |

## Supplementary Figure Legends

**Fig. S1.**
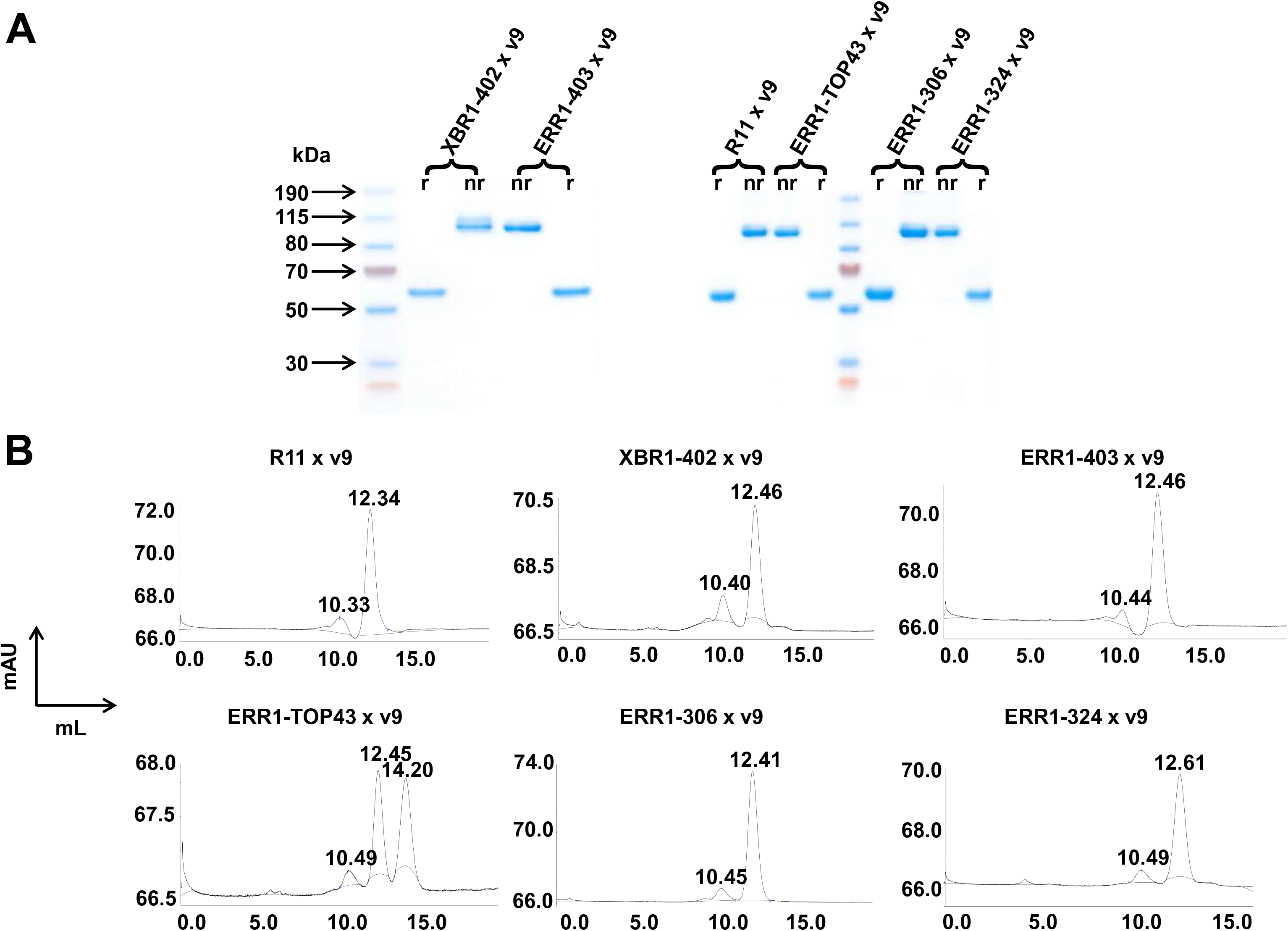
Generation of a panel of bispecific ROR1 × CD3 scFv-Fc. (A) SDS-PAGE and Coomassie Blue staining analysis of purified bispecific scFv-Fc showing the expected bands at ~100 kDa under nonreducing (nr) and ~50 kDa under reducing (r) conditions. (**B**) SEC analysis of bispecific scFv-Fc eluting as major peak at ~12.5 mL. The ~10.4-mL minor higher molecular weight peaks indicate the level of aggregation. A 14.20-mL lower molecular weight contamination in the ERR1-TOP43 × v9 preparation was not detectable by SDS-PAGE and Coomassie Blue staining.

**Fig. S2.**
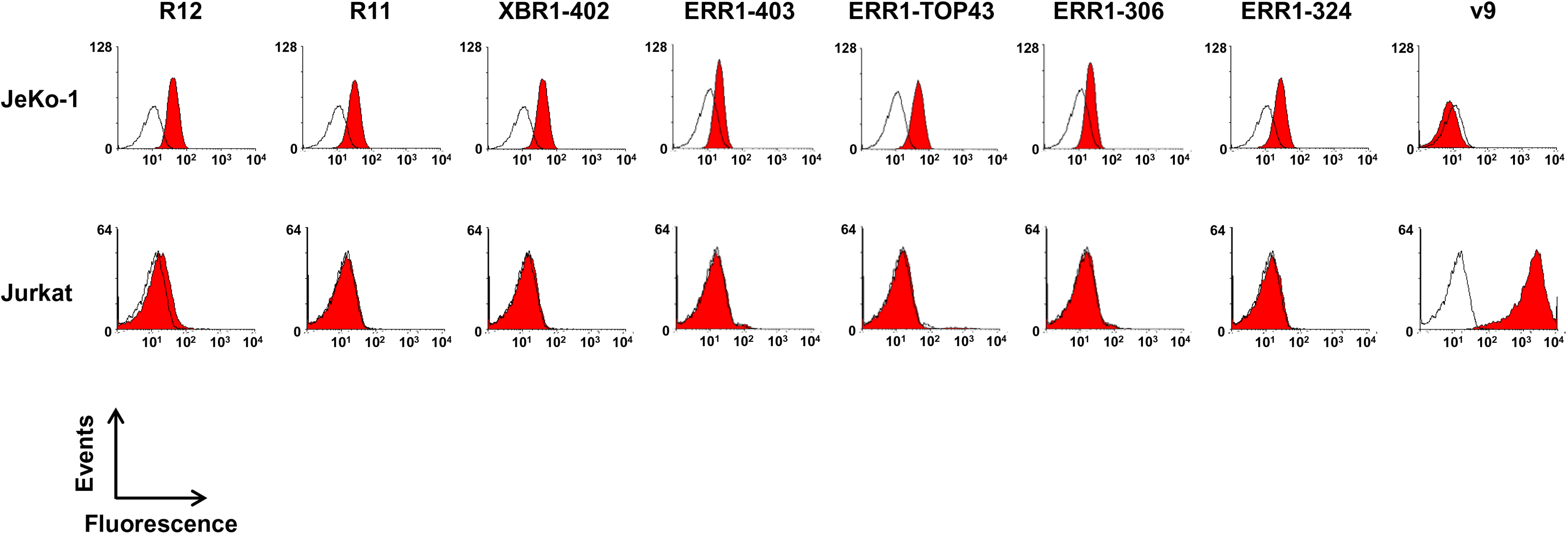
T-cell engagement mediated by bispecific ROR1 × CD3 scFv-Fc. The indicated monospecific scFv-Fc were analyzed for binding to human ROR1-positive cell line JeKo-1 and human CD3-positive T-cell line Jurkat by flow cytometry at a concentration of 5 μg/mL followed by Alexa Fluor 647-conjugated goat anti-human IgG pAbs. Open histograms show the background binding signal of the secondary pAbs.

**Fig. S3.**
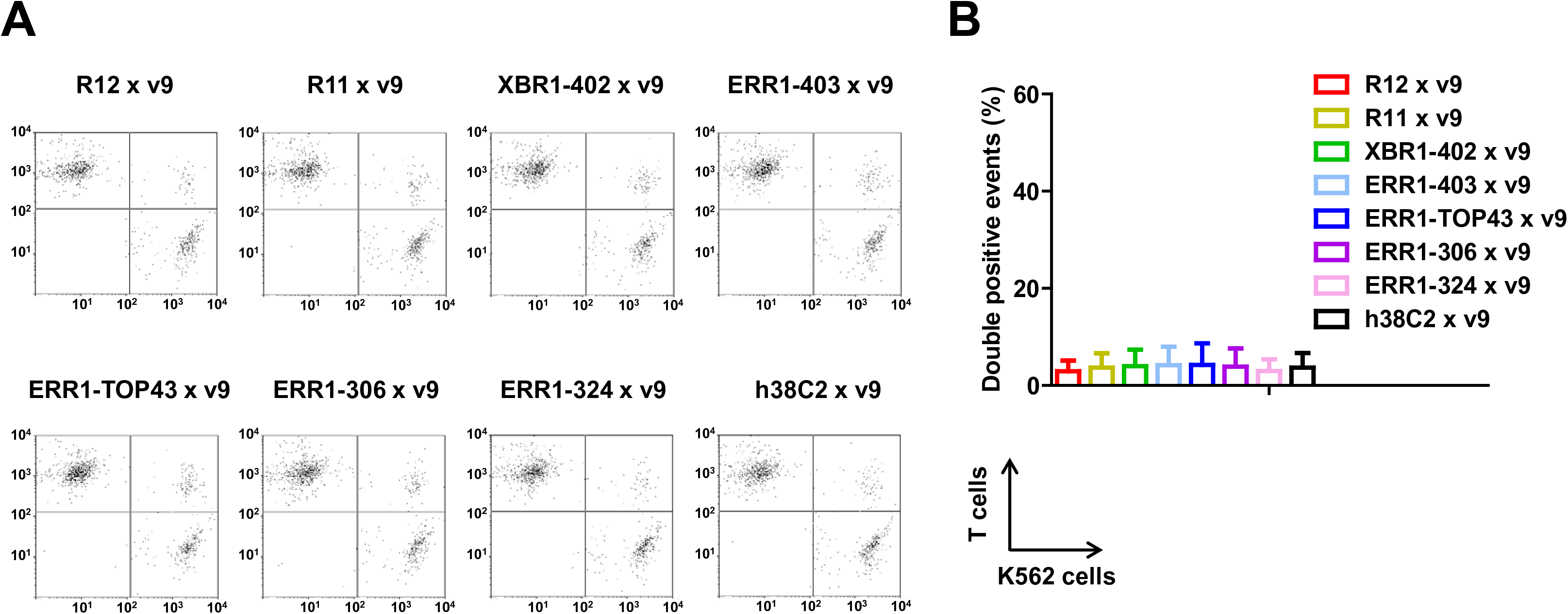
Cell surface binding of monospecific ROR1 and CD3 scFv-Fc. (**A**) Crosslinking of 5 × 10^4^ primary human T cells (stained with CellTrace Far Red) and 5 × 10^4^ K562 cells (stained with CellTrace CFSE) in the presence of 1 µg/mL ROR1 × CD3 and control scFv-Fc. Double stained events were detected by flow cytometry. (**B**) Quantification of double stained events from 3 independent triplicates (mean ± SD).

**Fig. S4.**
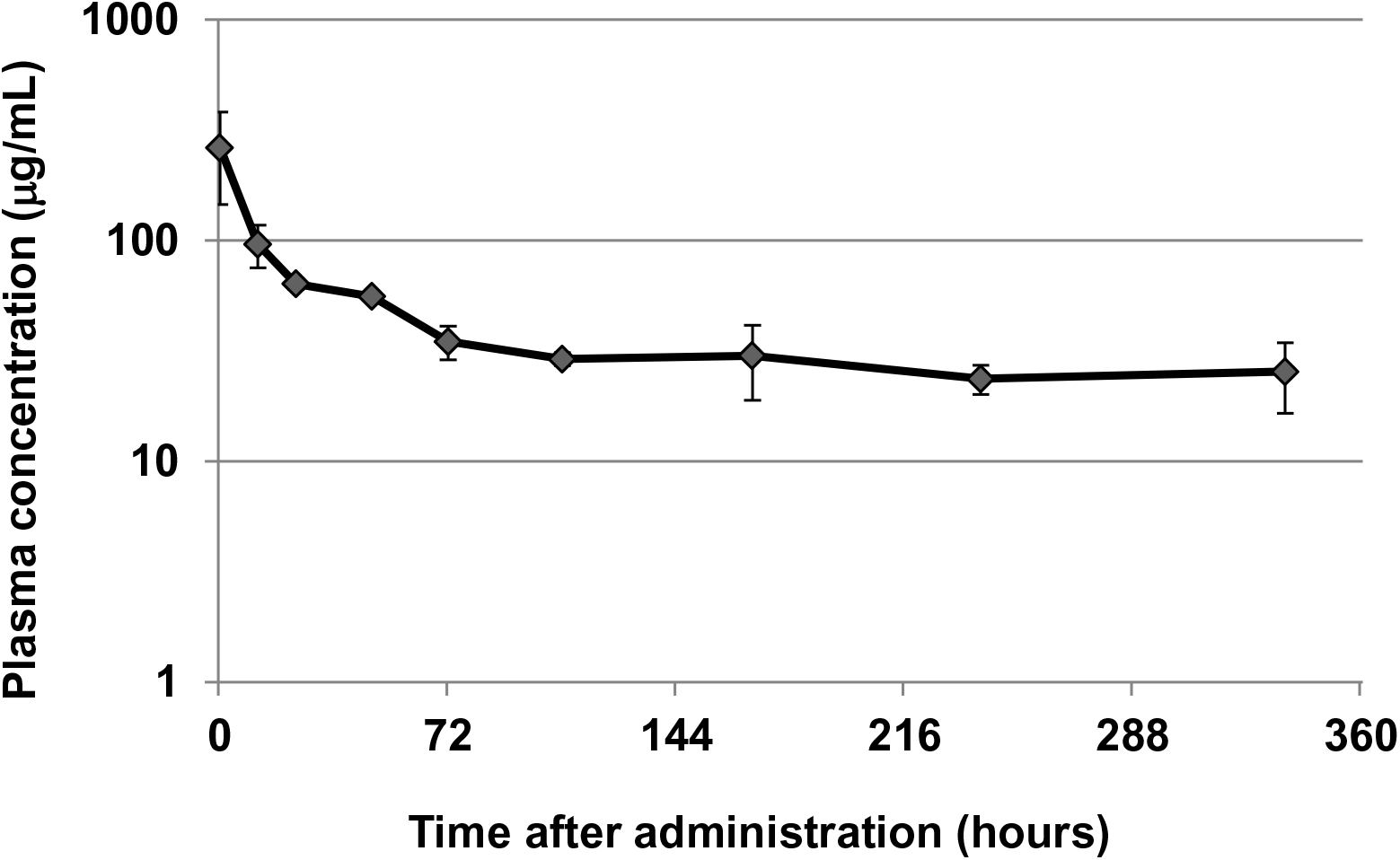
Pharmacokinetics of R11 × v9 scFv-Fc. Five female CD-1 mice were injected i.v. with 6 mg/kg R11 × v9 scFv-Fc. The plasma concentrations of the biAb at the indicated time points were quantified with a sandwich ELISA, using ROR1 ECD for capture and peroxidase-conjugated goat anti-human IgG pAbs for detection. Shown are mean ± SD values for each time point. The pharmacokinetic parameters are listed in Supplementary Table S2.

**Fig. S5.**
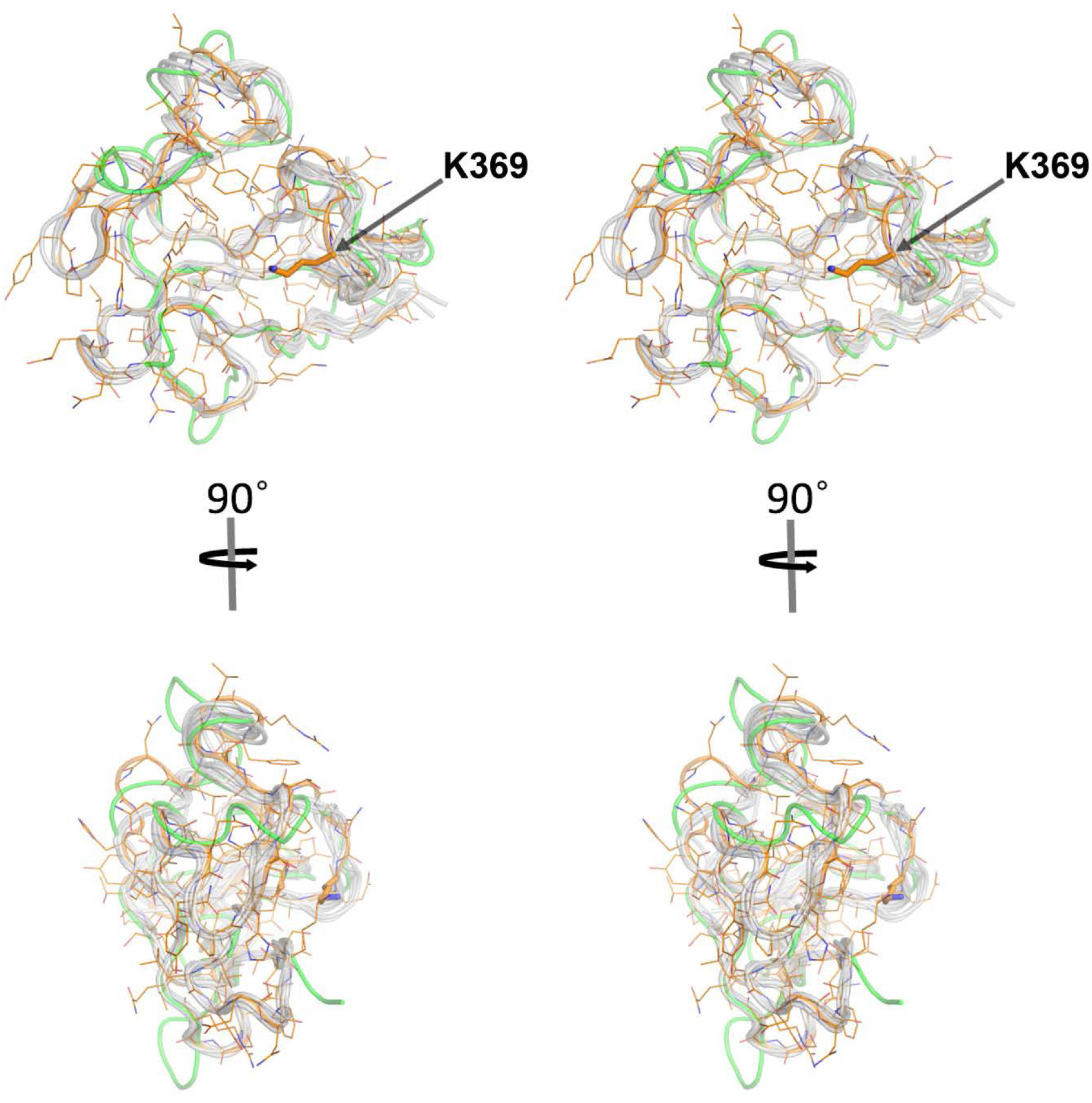
Superposition of the Kr domain of ROR1 with other crystallized kringle domains. Stereo images of the Kr domains of human ROR1 (orange; PBD: 6BA5), human plasminogen (grey; PBD: 1KRN, 1PMK, 1CEA, and 1KIO), human ApoA (grey; PBD: 4BVD), human hepatocyte growth factor (grey PBD: 5COE), and human Kremen1 (green; PBD: 5FWW). The canonical surface pocket of the kringle domain is constricted in the Kr domain of ROR1 due to an inward conformation of loop 5 and due to partial occupation by the side chain of Lys369 (arrow).

**Fig. S6.**
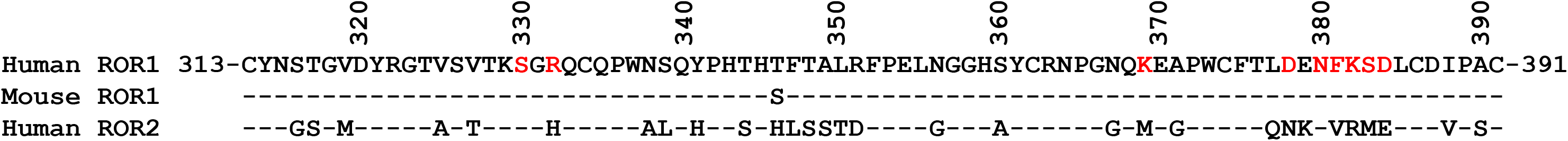
Amino acid sequence alignment of the Kr domains of human ROR1, mouse ROR1, and human ROR2. Mouse ROR1 (77/78; 99 %) and human ROR2 positions (48/78; 62 %) that are identical to human ROR1 are shown as dashes. Amino acid residues forming the epitope of scFv R11 are marked in red.

